# Non-ergodicity in ecology and evolution

**DOI:** 10.1101/2025.09.01.673433

**Authors:** Teemu Kuosmanen, Alexandre Minetto, Ville Mustonen

## Abstract

Stochasticity plays an important role in all biological systems. The standard way to deal with stochasticity involves averaging over an ensemble of independent realizations. However, such mean statistics need not accurately reflect the typical outcomes in any finite sample unless the system satisfies the property of ergodicity, which guarantees that each trajectory will over time experience the same statistics as the entire ensemble. Here, we argue that, in contrast, *non-ergodicity* might instead be the rule rather than exception in real biological systems and investigate its implications for eco-evolutionary dynamics through three case-studies. First, we show how demographic stochasticity leads to ergodicity breaking where the asymptotic growth rate carries a signature of the initial condition. This motivates us to define a mutant establishment threshold, which quantifies a critical population size above which the typical mutant population starts to grow. Second, we consider environmental stochasticity and demonstrate that autocorrelated environmental variation does not average out in time, which has the important consequence that the fitness of a genotype cannot be simply averaged over the environments. Finally, we show how in a metapopulation structure the evolutionary dynamics within a typical subpopulation can deviate from the ensemble dynamics in the entire metapopulation, which is sufficient to explain the evolution and persistence of cooperation despite a fitness cost.

## Introduction

Different types and levels of stochasticity invariably affect biological systems on all scales, from the molecular biology of a single cell to the ecology and evolution of populations, species, and ecosystems. Consequently, for organisms to function reliably, they must realise adaptation strategies that allow for robust regulatory control against environmental perturbations [1]. Demographic and environmental stochasticity can cause extinction of populations [2, 3] as well as shape the evolution of growth strategies [4], including reversing deterministic selection [5], reducing population turnover [6], and encouraging bet-hedging [7, 8]. All selectable variation is ultimately generated by mutational processes characterized by their random and undirected nature, but rapidly evolving cell populations can also leverage such stochasticity to adapt to stressful environments by increasing their mutation rate [9–11], an example of anti-fragility [12]. Indeed, it seems that stochasticity is not just “noise” that living systems must overcome, but rather an essential feature of the operating algorithm of both cellular function and evolution itself [13].

The standard way to deal with stochasticity generally involves averaging over an ensemble of repeated experiments in some way. Mathematically, the most tractable approach is often to focus on expected outcomes, defined as the average over infinitely many independent stochastic realizations, a quantity that paradigmatically serves as the object of maximization in various fields of science, including statistics, machine learning, economics, and decision-theory. However, it is well-known that such mean statistics need not accurately reflect the observed typical behaviour in any finite sample of outcomes. Indeed, the stochasticity related to any particular trajectory can be safely “averaged out” only in the case where the underlying system satisfies the property of ergodicity [14], which guarantees that all trajectories will eventually be statistically indistinguishable from the behaviour described by the corresponding mean-field theory.

While the ergodic assumption has been arguably useful in the context of thermodynamics and other effective physical theories, more complex systems often exhibit ergodicity breaking where the typical behaviour of individual trajectories deviates from the ensemble average. A canonical example of non-ergodicity is given by geometric Brownian motion (GBM), commonly employed for modelling asset prices [15], where the ensemble average grows exponentially indefinitely even though every trajectory is guaranteed to go ultimately ex-tinct due to the presence of multiplicative noise [16]. More recently, the idea of ergodicity breaking and its potential implications has been discussed in many other areas ranging from economics [17], human decision-making [18], medical and behavioural sciences [14, 19, 20], reinforcement learning [21], cell transport [22, 23], movement ecology [24, 25], and also evolution [26–28].

As part of the difficult task of defining the suitable fitness proxy in different types of models, eco-evolutionary theory is in principle familiar with the different types of averages, and the important distinction between arithmetic and geometric growth rates, which underlies the ergodicity breaking in GBM. However, we believe that the more general concept of non-ergodicity remains unfamiliar in the biological context even though ergodicity is com-monly assumed, either explicitly (e.g., [29]) or implicitly (e.g., [30]). Here we go beyond the much-studied GBM model and show how non-ergodicity arises in three distinct biologically motivated case-studies.

## Results

Consider a stochastic growth process *x* from which one can obtain sample trajectories (*x*_*i*_(0), …, *x*_*i*_(*t*)) as a function of time *t*. The asymptotic growth rate (or long-term average geometric growth rate)

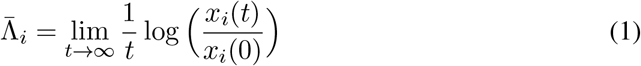

is one of the most widely used fitness concepts for simple growth processes. We use the overbar to emphasize that it is in fact a time-average, and it can be interpreted geometrically as the backwards extrapolated slope that provides the best linear fit for the growth trajectory on the log-scale. However, if the process is stochastic, 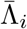 can also become stochastic, which motivates us to define 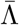 formally as the average over the eventually realized outcomes

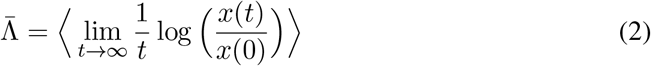

where, critically, the time limit is taken first before averaging. One of its key desirable properties is that a strain (or strategy) maximizing 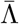 can be seen as the evolutionarily stable strategy (ESS) in the sense that it will almost surely outnumber any other strategy in the long run [31]. However, computing the time limit is generally mathematically untractable; in practice, one has to always estimate 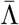 from a finite sample of *L* independent empirical realizations with finite time horizon as

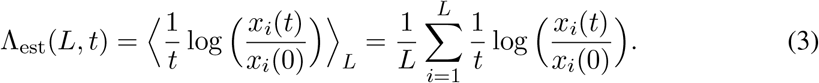

This estimate can be improved by either letting the process run longer or by increasing the number of trajectories in the ensemble, but ultimately it is the empirically realized fitness that one aims to characterize and predict.

Mathematically, if one knows the solution of the process *p*(*x, t*), one can formally obtain an analytical expression for Λ in the ensemble limit

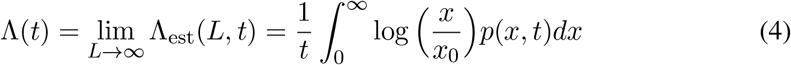

and then investigate the convergence to a theoretical value of Λ in the limit *t → ∞*. For example, for geometric Brownian motion defined by the stochastic differential equation (SDE)

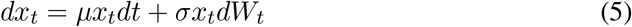

where *W*_*txg*_ is the standard Wiener process, it is straightforward to compute that Λ = *µ − σ*^2^*/*2 (see SI material), which can be recognized as the geometric growth rate of the process [16].

Importantly, however, Λ was derived by first taking the ensemble limit and only then the time limit, whereas in the definition of 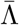 the time limit was taken first. If the process is ergodic, one can safely exchange the order of the limit operations such that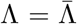, but generally, if the process is non-ergodic, this is not guaranteed. Biologically, the question of (non)ergodicity is then actually related to estimation: what is the possible error that we make if we try to predict the true empirical fitness with the theoretical value Λ?

This motivates us to consider the *bias* of the estimator

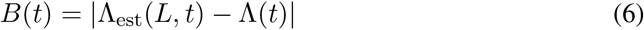

as a function of time and ensemble size. In particular, does the theoretical value converge to the empirical one in the sense that

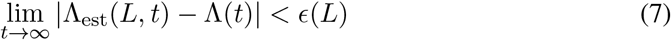

with *ϵ*(*L*) *→* 0 with *L → ∞*? Even if the estimator is unbiased in this sense, it remains important to verify the rate at which the bias decays to identify the timescale after which the ergodic approximation becomes valid [32].

The above approach makes the question of non-ergodicity concrete in terms of the exchangeability of the limit operations, but it only works if the ensemble average can be computed analytically. More generally, for any observable *f*_*i*_(*t*) of the process, one can consider a purely empirical metric

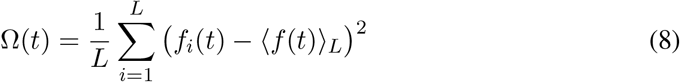

known as the Thirumalai-Mountain effective ergodicity metric [33], but it can also be inter-preted as the *variance* of the estimator. Intuitively, if the process is ergodic, every trajectory *i* will eventually be indistinguishable from the ensemble average, and thus Ω measures the degree of self-averaging, with ergodic processes having the property that Ω(*t*) *→* 0 with *t → ∞*. It then follows that if the process is non-ergodic in this sense, even an unbiased estimator will make an error that cannot be removed even in the limit of infinite data.

In what follows, we use these two metrics to quantify the non-ergodicity first, in two analytically tractable models for demographic and environmental stochasticity, and then in a metapopulation model. This allows us to show when and in what sense stochasticity averages out in eco-evolutionary dynamics. Furthermore, by clearly distinguishing between the ensemble average from the typical outcome, we show how both of them can be biologically meaningful and appropriate descriptors in different situations.

### Case 1: Demographic stochasticity

Although a useful starting point, GBM is not directly applicable to model the growth of biological populations, because the signal-to-noise ratio in the GBM remains constant, while it is widely established that demographic stochasticity should “average out” in large populations. But in what sense is this precisely true?

When the population dynamics is derived explicitly from the underlying birth and death rates [6, 34, 35], a SDE of the following form is recovered

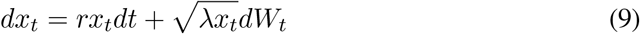

where *x*_*t*_ is the population size, *r* := *β−δ* gives the expected per capita growth rate (difference of the birth and death rates), and the demographic stochasticity scales with the square root of the population size and the population turnover rate *λ* := *β* +*δ* (see [6]). In the case where the birth and death rates remain constant, this equation reduces to Fellerian diffusion [36] which has a closed-form solution for the probability density *p*(*x, t*) with the absorbing boundary set at *x* = 0 to account for stochastic extinction (see Methods for more details).

We can now compute Λ for this process by taking the time limit of [4], which yields

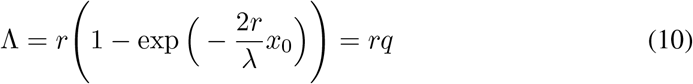

where *q* can be interpreted as the ultimate establishment probability (see Methods). Intriguingly, this result shows how the asymptotic growth rate Λ now explicitly depends on the initial condition *x*_0_, which can be considered a hallmark of non-ergodicity (see also [37] where this was empirically observed). We verify this using stochastic simulations (see Fig. 1B), which confirm that there exists a critical population size, which depends on the growth parameters *r* and *λ*, below which Λ is sensitive to the initial condition while above it Λ remains close to the ensemble average *r*. This is in sharp contrast to GBM, where the asymptotic growth rate does not depend on the initial condition at all.

**Figure 1.**
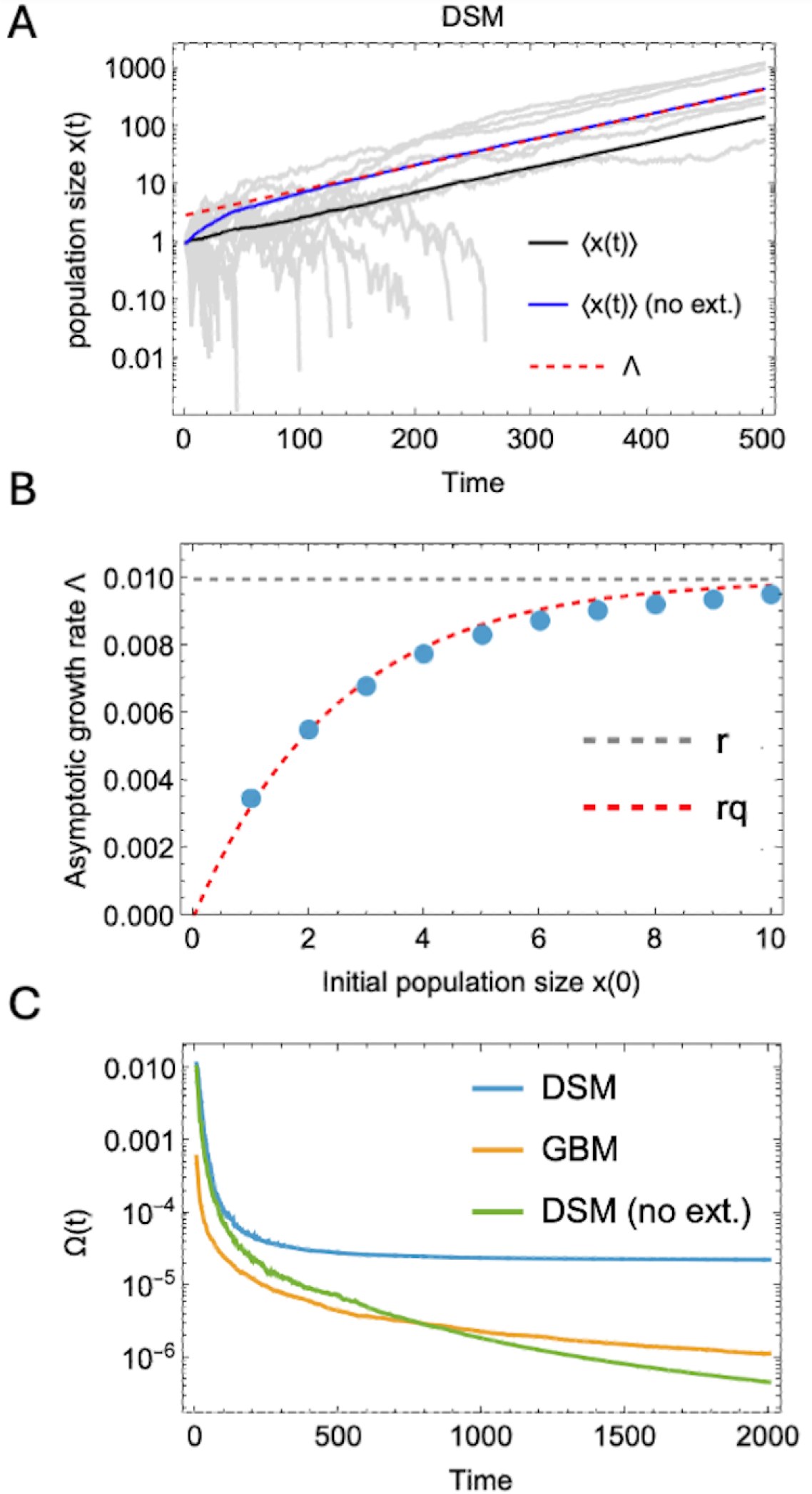
on-ergodicity in demographic stochasticity models. **A** Example trajectories of the demographic stochasticity model (DSM) defined by [9] shown in gray; ensemble average of the process grows exponentially (black line). If conditioned on non-extinction, the ensemble grows initially superexponentially (blue line) before settling to grow with the expected growth rate *r*. If extrapolated backwards in time, it appears as if the growth had started approximately from the extinction threshold 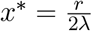 instead of the true initial condition *x*_0_. **B** The asymptotic growth rate Λ is computed by first taking the time-limit before averaging. Stochastic simulations starting from different initial conditions *x*_0_ reveal that Λ depends on the initial condition as correctly predicted by [10]. **C** Quantifying non-ergodicity with the Thirumalai-Mountain metric Ω(*t*) which can be interpreted as the variance of the Λ-estimator. The DSM model is even more non-ergodic than GBM; however, conditioning on non-extinction restores ergodicity. Parameters: *r* = 0.01, *λ* = 0.05, *σ* = 0.05 end-time: *T* = 2000, no. trajectories = 1000.

We then further compared the demographic stochasticity model [9] with GBM by computing the bias and variance of the Λ-estimator. Fig. 1C shows that the variance of the demographic stochasticity model is higher than in the GBM and converges to a positive value unlike in the GBM, where it continues to decay. Thus, the demographic stochasticity model is actually even more non-ergodic than GBM as measured by the Thirumalai-Mountain metric. However, if one conditions the process [9] with non-extinction, the variance quickly decays to zero, implying that the non-ergodicity arises mainly due to presence of the extinction boundary. This is, however, a biologically important property of the model, because all mutations must arise from a single individual, which has a significant risk of stochastic extinction [38].

To characterize the apparent stochastic extinction boundary more precisely, we turn to analyze the log-transformed process

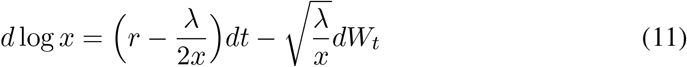

which can be obtained by applying Itô’s Lemma to [9]. We notice that the deterministic part of [11] explicitly depends on the current population size such that it is initially negative and switches to become positive at the critical size

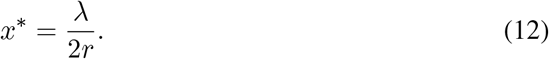

We call this the extinction threshold, which allows one to better understand the early stochastic phase of growth that the ensemble average is hiding. Namely, if the population size falls below this value, it’s expected log-fitness becomes negative and the typical trajectory goes extinct despite having a positive deterministic growth rate. We emphasize that this threshold is not “hard” in the sense of an Allee threshold, where the deterministic growth rate becomes negative, but it only describes the fate of a typical trajectory. Also, because the process is Markovian, the typical outcome can change if the trajectory crosses the boundary at any point.

We further note that if we condition the process on non-extinction, the ensemble average can be written as

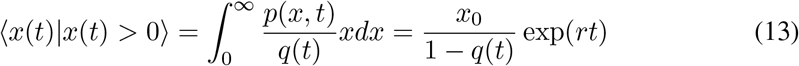

and when extrapolated backwards in time, it appears as if the exponential growth had started from the initial size

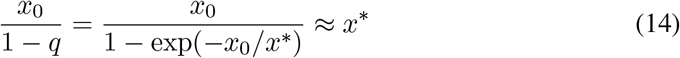

that is, the extinction threshold is in fact the first-order approximation for the effective onset of the exponential growth (see Fig. 1A).

This concept can be readily generalized to more complicated situations. Consider now a mutant-resident model where two types *x*_1_ and *x*_2_ are competing following their own dynamics according to [9]. Kuosmanen *et al*. [6] show that the mutant frequency *p*_1_ = *x*_1_*/*(*x*_1_ + *x*_2_) then follows the SDE

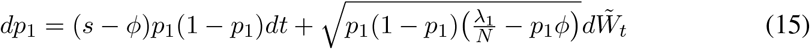

where *s* = *r*_1_ *− r*_2_ is the selection coefficient (difference of the per capita growth rates), the total population size *N* = *x*_1_ + *x*_2_ grows approximately deterministically with the mean fitness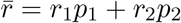, and 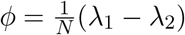 is the so-called turnover flux which becomes negligible in large populations. We can now similarly solve the mutant *establishment threshold* from the log-transformed equation of [15] as

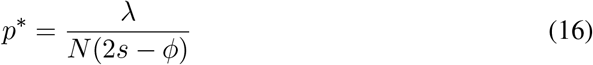

which gives the frequency above which the prevalence of the typical mutant under positive selection starts to grow and dominates genetic drift. Indeed, in the Wright-Fisher limit of constant turnover (*λ*_1_, *λ*_2_) *→* (1, 1), the establishment threshold simplifies to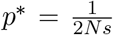, which appears in the existing population genetics literature [39–41] as the approximate onset of the exponential growth phase where the strength of deterministic selection overcomes the stochastic forces of genetic drift.

Here we extend these observations by showing how the establishment threshold depends on the turnover rates and has a natural interpretation by describing the typical mutant dynamics (see Fig. 2). If the mutant population starts to grow below the threshold, the typical mutant trajectory goes extinct since the expected log-frequency change is negative. However, if the mutant manages to temporarily cross this threshold by aid of stochastic fluctuations, this prediction is updated, and the fate of a typical mutant is to establish as part of the population, and thus avoid immediate extinction. Single trajectories can naturally cross the boundary multiple times, and each time the qualitative prediction is updated accordingly.

**Figure 2.**
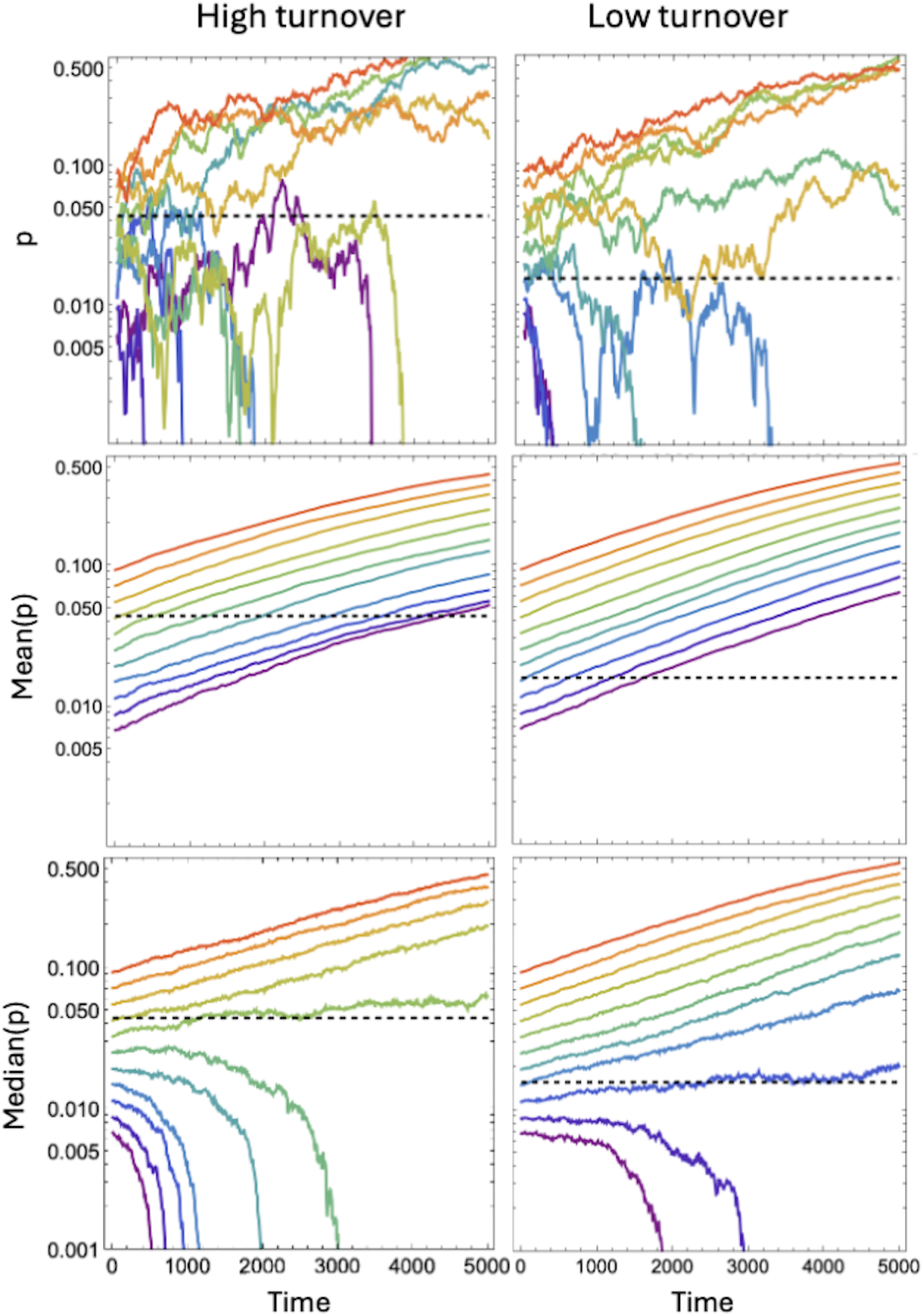
Establishment threshold predicts the fate of a typical mutant. Evolutionary dynamics of a mutant under positive selection starting from different initial frequencies *p*(0). **First row:** Examples of individual evolutionary trajectories of the mutant frequency *p*(*t*) as a function of time. The high-turnover mutant (left column) experiences higher fluctuations than the low-turnover mutant (right column) and is more likely to go extinct. **Second row:** The mean mutant frequency *⟨p*(*t*)*⟩* grows exponentially irrespective of the initial condition or the turnover rate. **Third row:** The median mutant frequencies on the other hand show clear initial condition dependence: mutants starting below the establishment threshold *p*^*∗*^ = *λ/*2*Ns* (dashed lines) typically go extinct while those starting above the threshold typically establish. The establishment threshold also depends on the turnover rate: mutants with lower turnover establish at a lower frequency. See Methods for more details.

### Case 2: Environmental stochasticity

We next consider environmental stochasticity where the birth and death rates themselves, and not just the timing of individual events, can fluctuate. Because of this, environmental stochasticity can affect also large populations [3]. However, it is commonly assumed that the effects of environmental stochasticity average out in time.

Mathematically, we can model environmental stochasticity with a system of SDEs, where the environment also has some dynamics, which in real systems typically also depends on the organismal impact leading to eco-evolutionary feedback:

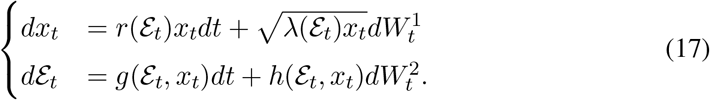

As we want to focus specifically on the effects of environmental stochasticity, we ignore demographic stochasticity for now and model the population growth as a deterministic process

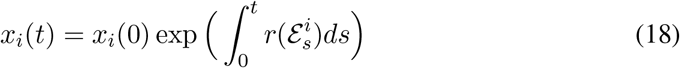

where the growth rate still depends on the particular sequence of environmental states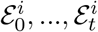, of the stochastic process *d*ℰ_*t*_. Now, assuming that the environment is ergodic, we can express the long-term growth rate [1] as

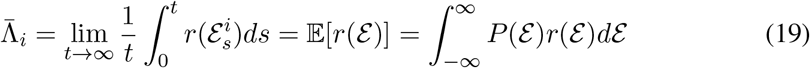

where we now explicitly made use of the ergodicity assumption, which allowed us to replace the time-average with an average over the probability space *P* (ℰ). In the simpler case, where the sequence of environments are independent and identically distributed, the law of large numbers can be applied to derive an analogous result [31]. The essence of the above result is that it allows one to change an explicitly open time problem to an integral over the space of environments with the important consequence that the environment, if ergodic, can be treated as a confounder that can be simply averaged over to determine the fitness. But how good is the ergodic assumption for biological systems? In particular, how should the probability distribution *P* (ℰ) be interpreted if it becomes coupled with the organismal impact (e.g., as in Eq. [17])? And how can fitness be defined for non-ergodic environments?

As a simple analytically tractable case study, let us begin by considering a family of Gaussian fitness functions of the form

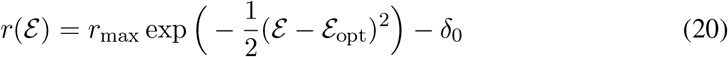

and assuming that the environment follows the Ornstein-Uhlenbeck process (for previous modelling, see e.g., [34])

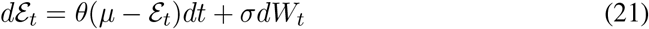

which is the only Markovian process admitting a Gaussian stationary distribution. Biologically, it can be interpreted as describing a model where the population attempts to equilibrate a diffusing environment around the value *µ*. In the limit *θ → ∞*, where this environmental regulation is essentially “perfect”, the process [21] reduces to white noise [42], which is the simplest form of environmental stochasticity also known as micro-evolutionary fitness seascape noise [43, 44] commonly employed in modelling. On the other hand, in the limit *θ →* 0, the process [21] becomes a pure diffusion.

We can compute the long-term growth rate for this case analytically from [19] as (see SI

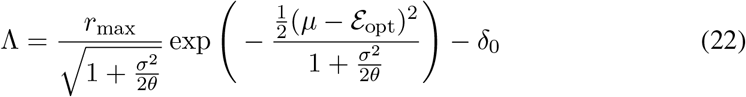

where *r*_max_ is the highest achievable growth rate at the environmental optimum 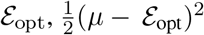 is the expected distance from the environmental optimum, *σ* is the diffusion constant of the environment, and *θ* is the regulation parameter that gives the characteristic timescale for the process to mean revert (all of which can be more generally trait specific parameters). We notice that in the white noise limit *θ → ∞*, the enviro nmental stochastic ity averages out in time leading to long-term growth rate 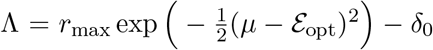 which is independent from the noise level *σ*, but only depends on the mismatch of the environmental statistics that the population can learn to infer [30]. On the other hand, in the limit of no regulation *θ →* 0, the environment will diffuse away from the habitable region and the population will decay with Λ = *−δ*_0_, further implying a threshold value for *θ* that is needed to allow the population persist.

In these intermediate values of *θ*, the process [21] is still autoregressive, but the noise becomes autocorrelated with states ℰ_*s*_ and ℰ_*t*_ having a covariance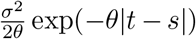. Indeed, autocorrelated environmental stochasticity is a notable feature of real ecosystems [45, 46], but it remains to be incorporated into models. Fig. 3 shows how such autocorrelated stochasticity introduces temporary non-ergodicity before the process eventually self-averages at a characteristic timescale set by the parameter *θ*.

**Figure 3.**
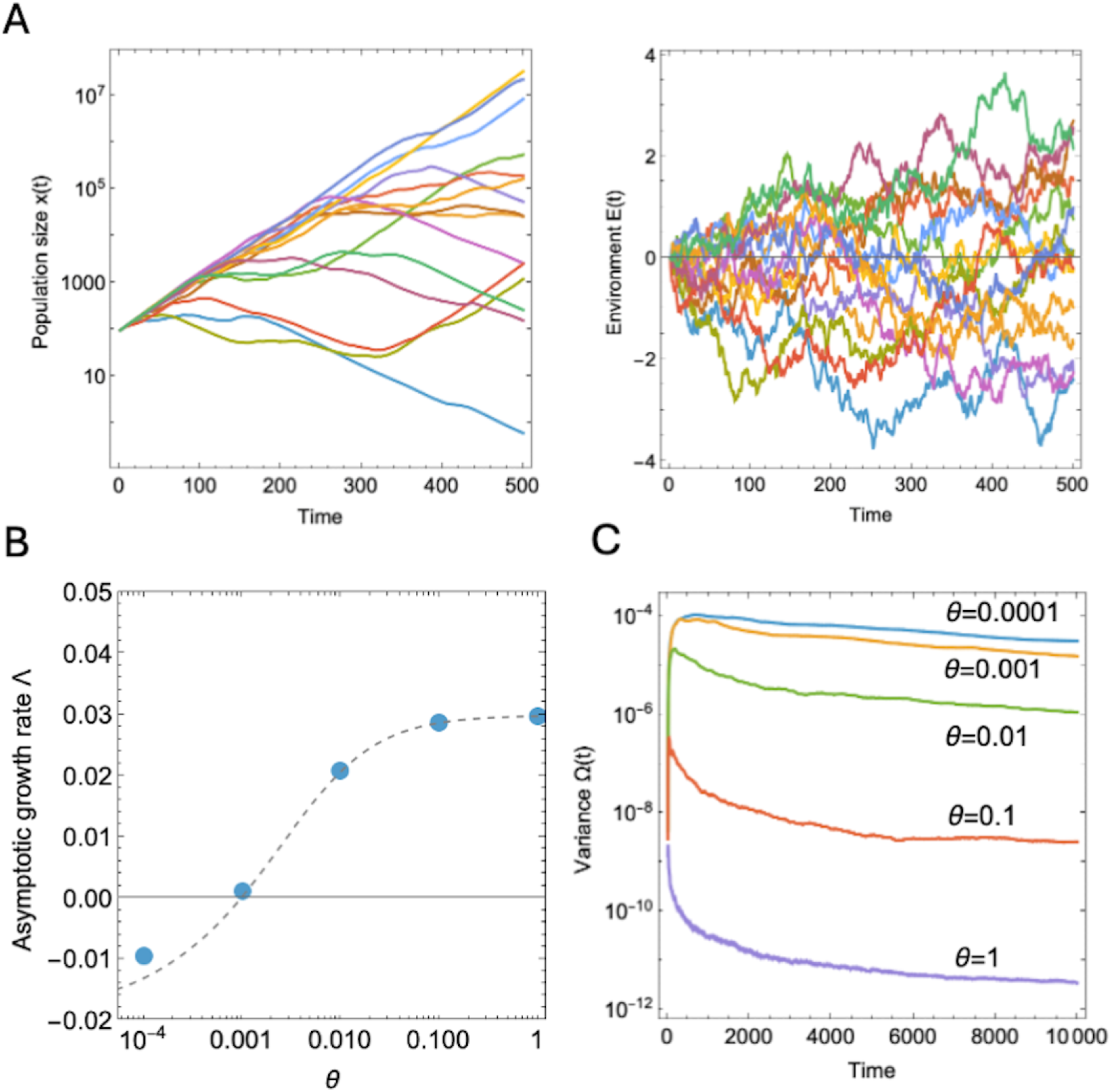
on-ergodicity in environmental stochasticity model. **A** Example population (left) and environment (right) trajectories of the environmental stochasticity model [20-21] with *θ* = 0.001. **B** The asymptotic growth rate Λ as a function of the parameter *θ* which sets the typical timescale of environmental autocorrelation. Analytical formula [22] (dashed line) holds, but convergence is very slow for small values of *θ*. **C** Corresponding values of the Ω-metric for different values of *θ* which shows that the environmental stochasticity model exhibits temporary non-ergodicity. Parameters: *µ* = 0, *λ* = 0 (no demographic stochasticity), *σ* = 0.1, ℰ_opt_ = 0, *r*_max_ = 0.05, *δ*_0_ = 0.02, *x*_0_ = 100, ℰ_0_ = 0, end-time: *T* = 10^4^, no. trajectories = 100.

In reality, however, also the environmental steady state can drift in time, i.e. *µ → µ*_*t*_, either due to organismal impact or exogeneous environmental processes. In this case, the process will no longer be autoregressive meaning that the trajectory statistics will no longer match the ensemble statistics, and consequently ergodicity will no longer be restored even in the long time limit, but instead the mismatch between ensemble and time-averages can continue to grow (see Fig. 4).

**Figure 4.**
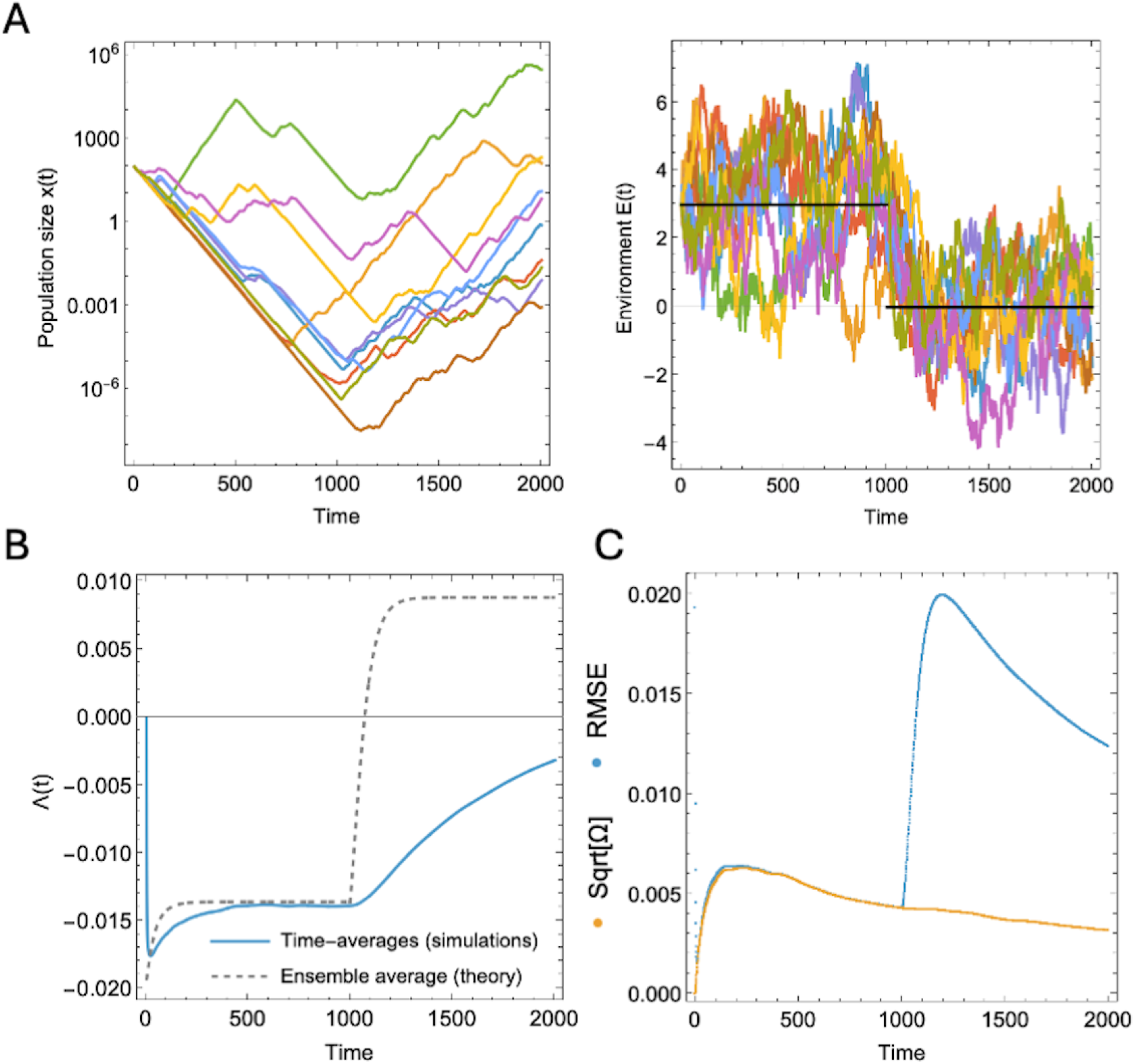
Λ-estimator becomes biased in non-autoregressive environments. **A** Example trajectories of the environmental stochasticity model with *θ* = 0.01 and a time-dependent environmental equilibrium switching from initial value *µ*_0_ = 3 to *µ*_*t*_ = 0 at time *t* = 1000. **B** When the environment is non-autoregressive, the environmental statistics along different trajectories can deviate from the ensemble statistics and the temporary non-ergodicity can lead to accumulating tracking error. **C** The root mean square error of the Λ-estimator increases even though the variance decreases. This implies that the ensemble average becomes increasingly biased.

### Case 3: Population structure

In previous case studies, we showed how demographic and environmental stochasticity can lead to non-ergodicity via the extinction boundary and autocorrelated fitness fluctuations. Next, we show how population structure can also lead to a different kind of ergodicity breaking by considering a metapopulation structure where both the ensemble average and typical subpopulation average have a clear biological meaning.

A metapopulation describes independent parallel population dynamics within spatially separated subpopulations connected by a possibility of rare dispersal events. Metapopulation structures arise naturally at different scales ranging from habitable island patches to entire planetary ecosystems. A characteristic feature of a metapopulation is the possible extinction of local subpopulations and the presence of empty patches, which can be critical for the overall success of the metapopulation by allowing reinvasion by later dispersal [47]. As we will next show, this mechanism can lead to non-ergodicity where the ensemble frequency in the entire metapopulation can diverge both quantitatively and qualitatively from the typical dynamics within any subpopulation.

Consider the well-known public good game [48], which provides a classical model for the evolution of cooperation, but now in a metapopulation context, where each of the *m* = 1, …, *M* subpopulations follow their own local public good dynamics. We consider the evolutionary dynamics of two genotypes, the cooperators (C) which contribute to the production of a public good at a fitness cost, and defectors (D) which do not, but nevertheless can enjoy access to the local public good, which is distributed equally to all the members of the subpopulation. This leads to frequency-dependent fitness functions:

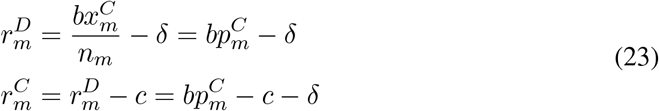

where 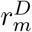 is the defector fitness in subpopulation *m* with cooperator frequency 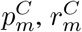, is the corresponding cooperator fitness, *b* is the growth-benefit of the public good, *c* is the individual-level cost of producing the public good, and *δ* is the death rate which determines the minimum amount of public good production that allows the subpopulation to persist. This leads to the following deterministic dynamics for the cooperator frequency *p*_*m*_ and total population size *n*_*m*_ of subpopulation *m*:

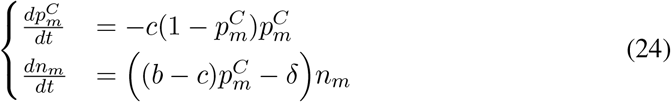

Thus, without invoking any other mechanisms that could maintain cooperation, the typical cooperator frequency will decrease according to the standard replicator equation in every subpopulation, which will eventually lead to the extinction of that subpopulation. But what ultimately matters for evolution is the metapopulation frequency of cooperators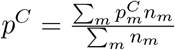, which critically must be computed as a weighted average of the subpopulation frequencies to give each individual equal weight. By simulating the public good dynamics with varying fitness costs *c* and metapopulation sizes *M*, and including the possibility for dispersal and mutation (see Methods for more details), we show how cooperation can be maintained in the metapopulation despite a significant fitness cost solely by virtue of the metapopulation structure (see Fig. 5).

**Figure 5.**
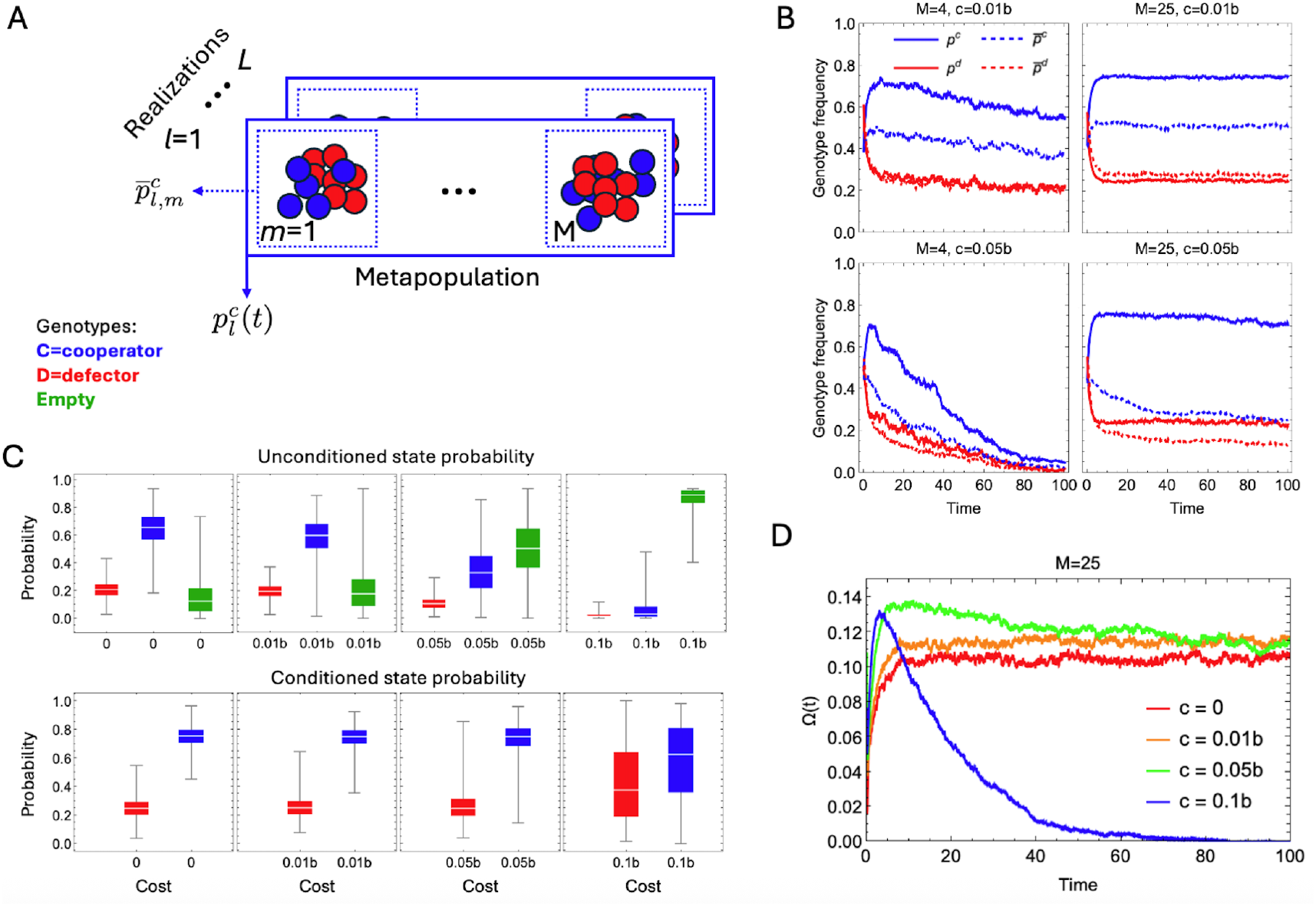
on-ergodicity in a metapopulation model. **A** Schematic of stochastic public good dynamics [23] simulated in a metapopulation setting. We define two types of biologically meaningful averages for cooperator and defector frequencies for each replicate *l* of a metacommunity *m* and time point *t*. First, 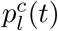 describes the global metapopulation frequency of cooperators (blue cells), 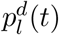 defectors (red cells). Second, 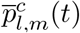 describes the per patch frequency of cooperators and similarly 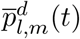for defectors. **B** The ensemble averages (solid lines) give the full metapopulation frequencies of cooperator (blue, *p*^*c*^) and defector (red, *p*^*d*^) genotypes, while the per patch-averaged frequencies (dashed lines, 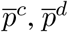,) represent typical genotype frequencies in subpopulations. Selection acts on the ensemble frequency, which is systematically higher than the typical cooperator frequency in any given subpopulation. With higher cost of cooperation (bottom row), the ensemble average approaches the typical frequency, but in larger metapopulations (second row) the ensemble frequency can nevertheless remain high even with a high fitness cost. **C** When the cost of cooperation increases, the state probabilities of cooperating subpopulations go down and the fraction of empty patches increase (top row). However, if conditioned on non-extinction, the state probabilities of cooperating subpopulations remain approximately the same suggesting that the conditioned dynamics is driven mainly by the absolute fitness effects. **D** Thirumalai-Mountain metric applied to the cooperator frequencies (see Methods for more details) reveals that the degree of non-ergodicity increases with the cost unless the metapopulation goes extinct which returns ergodicity.

This result can be understood mathematically by deriving a generalized replicator dynamics for the metapopulation frequency that takes into account how the population sizes of the subpopulations, which act as weights for the subpopulation frequencies, change over time. Let us denote 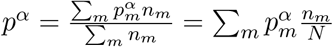 the metapopulation frequency of type *α*, where *N* is now the total population size of the metapopulation. We get that

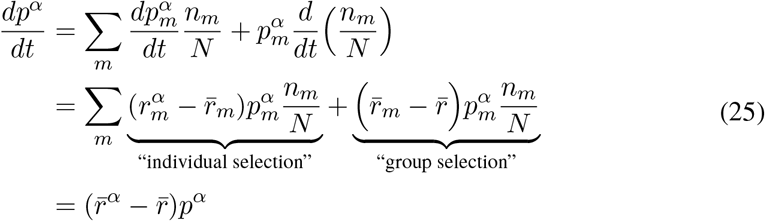

where we defined 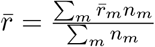 as the mean growth rate of the metapopulation, and 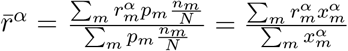 as the average growth rate of type *α* (*C* or *D*) weighted by their subpopulation sizes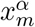. This result shows how also the metapopulation frequency follows a more general form of replicator dynamics, which can be formally decomposed to standard selection acting independently at each subpopulation, and “group selection” acting at the level of absolute fitness between the groups. Thus, even if the frequency of cooperators decreases in every subpopulation, the total metapopulation frequency of cooperators can still increase because highly cooperating subpopulations tend to outgrow defector-dominated subpopulations. This curious phenomenon represents a well-known example of a Simpson’s paradox, which has also been empirically observed [49].

We note that when the more general result [25] is applied to the public good dynamics [24], the condition for positive selection of cooperators can be written equivalently as

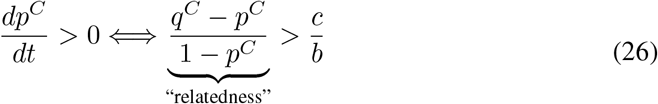

where 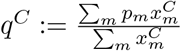 is another type of weighted cooperator frequency. This can readily be interpreted as Hamilton’s rule, because the left-hand side measures a type of relatedness, that is, the excess probability of sharing the cooperating genotype above the population average *p*^*C*^. However, in contrast to the generalized replicator dynamics from which it was derived, the Hamilton’s rule only gives the correct sign of selection and it cannot be used to integrate the full dynamics correctly. For example, one could expect based on the Hamilton’s rule alone that increasing the fitness cost of public good production would decrease the level of cooperation. Remarkably, this is not what happens in large enough metapopulations, which instead show surprisingly robust cooperation despite a significant fitness cost (see Fig. 5). This is because the correct relatedness measure actually depends implicitly on both parameters *b* and *c*.

Fig. 5C shows that while the fraction of empty patches increases with *c*, the persistence times of cooperating subpopulations remain the same when conditioned on non-extinction.

Thus, the conditioned dynamics is essentially independent of the fitness cost of cooperation, but is rather determined by the absolute fitness effects. The evolution of cooperation appears paradoxical because, in any given subpopulation, it typically is. However, when one appreciates that the ensemble behaviour can deviate from typical outcomes, the evolution of cooperation can readily make sense in light of non-ergodicity. Fig. 5D shows that the evolutionary dynamics of cooperation becomes increasingly non-ergodic with increasing fitness cost *c* according to the Thirumalai-Mountain metric applied to the cooperator frequency, unless the entire metapopulation goes extinct, which restores ergodicity.

## Discussion

Ergodicity is a powerful mathematical concept that can greatly facilitate the analytical treatment of complex systems by allowing to freely exchange averaging of single trajectories over time with ensemble averages over repeated experiments. This implies that ergodic systems are conservative, path-independent, and eventually forget their initial state. In contrast, biological evolution is widely regarded as open-ended, cumulative, and thus path-dependent, dissipative, and operating in states that are increasingly atypical and far from the thermodynamical equilibrium. Yet, much of the existing eco-evolutionary theory makes use of the ergodic assumption either implicitly or explicitly, and with few notable exceptions [19, 26, 28, 50], the concept itself seems not to be widely established in this context.

Here we have attempted to demonstrate through three concrete and biologically motivated case studies how ergodicity is directly linked to some of the most fundamental questions in eco-evolutionary theory and what the potential implications of non-ergodicity might be. First, we moved beyond the much-studied GBM and investigated two analytically tractable stochastic models that showcase the effects of demographic and environmental stochasticity. By computing the long-term growth rate Λ analytically, we showed how the extinction boundary and autocorrelated fitness fluctuations can lead to non-ergodicity, and characterized precisely in what sense demographic and environmental stochasticity can and cannot be averaged out. Then, we investigated evolution of cooperation in a metapopulation model, and showed how this classical problem can be reinterpreted in light of non-ergodicity.

Even if the system of interest is strictly speaking non-ergodic, it remains important to quantify the ramifications of using the ergodic approximation to describe the underlying non-ergodic behaviour. On the other hand, even if ergodicity is restored in the stationary limit, the temporary non-ergodicity in the non-stationary phase can have biologically important consequences as well. We approached this by systematically investigating how the bias and variance of the Λ-estimator behave as a function of time and ensemble size. In more complicated cases, where the process cannot be described analytically, ergodicity can be treated as a statistical phenomenon which can be inferred from time series data to which end several different estimators have been developed. Here we used the Thirumalai-Mountain effective ergodicity metric [33], which measures the degree of self-averaging over time and essentially corresponds to the (sample) variance of the estimator, which can be computed empirically even if the true mean value is not known.

We summarize our main findings as follows: (i) demographic stochasticity in the presence of an extinction boundary leads to non-ergodicity, but conditioning on non-extinction restores ergodicity; (ii) environmental stochasticity need not average out if the environmental fluctuations become correlated or coupled to population dynamics, which can lead to ergodicity breaking; (iii) non-ergodicity can manifest also at the level of population structure, where the typical evolutionary dynamics can qualitatively differ from the ensemble dynamics: the longterm growth rate of a metapopulation can exceed the average growth rate of its constituent subpopulations (this is analogous to finance, where a stock market index can systematically outperform the average returns of the underlying stocks it is composed of); (iv) taken together, non-ergodicity renders the notion of “fitness” even more obscure and challenges predictability because evolution becomes “computationally irreducible”; one cannot eliminate time from the equation, but must take the time-limit seriously; (v) yet, suitable conditioning can restore ergodicity and allow for normative predictions.

Our results show that the concept of ergodicity is also closely related to typicality, which has recently been proposed as an alternative normative theory in ecology and evolution [51]. By carefully distinguishing between the two ways of averaging without assuming ergodicity, new theory for the typical dynamics can be developed. We demonstrated this by developing the concept of establishment threshold, which better describes the behaviour of a single typical trajectory in contrast to the ensemble average, which hides such intuitive phenomena. Similarly, we showed how the typical dynamics within any given subpopulation can drastically deviate from the metapopulation dynamics. We believe that further investigations into typicality might prove to be highly useful: for example, when attempting to predict rapidly evolving pathogens, the ability to successfully establish in a typical invasion might be more predictive as opposed to anticipating the emergence of the fittest strain as the most likely adaptation [52].

## Materials and Methods

### Solution of the SDE

The analytical solution for the process [9] with the absorbing boundary set at *x* = 0 can be obtained by first solving the driftless process by expressing it as a Bessel process, and then using suitable time-transformations to account for the deterministic growth [53]. The solution of the driftless process with *r* = 0 and initial condition *x*(0) = *x*_0_ is given by

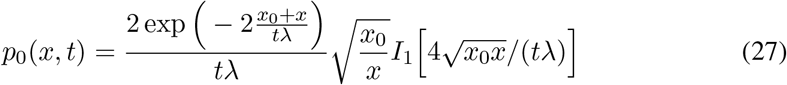

where *I*_1_ is the modified Bessel function of the first kind. Now, by transforming time as 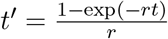, we can find the solution of the full process with drift as

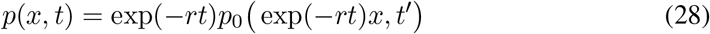

which can be easily verified as the solution of [9] by checking that it satisfies the associated Fokker-Planck equation (see Supplementary Material for more details).

One can now solve the fraction of non-extinct populations as a function of time as [36]

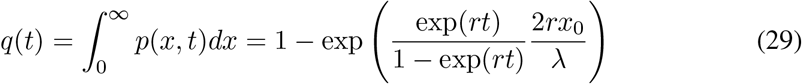

which further gives the ultimate establishment probability as

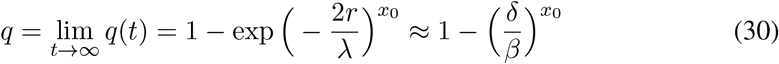

closely matching to the analogous branching process limit [7].

### Establishment threshold

Consider a resident population with birth and death rates *b*_0_ and *d*_0_, respectively, and a mutant with birth and death rates *b*_1_ and *d*_1_. In Fig. 2 we simulated the population dynamics using the patch model derived in Ref. [6], which equilibrates the resident population to size

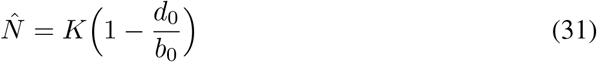

while allowing an approximately constant invasion fitness *s* = (*b*_1_ *− d*_1_) *−* (*b*_0_ *− d*_0_) for the mutant. Consequently, the establishment threshold [16] for the mutant is now approximately

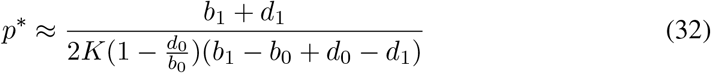

ignoring the turnover flux term which is of order *K*^*−*2^. We fix the resident strategy and consider two variants with equal strength of selection set to *Ks* = 10 but with varying turnovers *λ*_high_ = 1.5 *· λ*_0_ (high turnover mutant) and *λ*_low_ = 0.5 *· λ*_0_ (low turnover mutant) to contrast the impact of the turnover rate. We then simulated 1000 trajectories from each initial frequency and computed the mean and median mutant trajectories.

### Metapopulation model

We simulated the birth-death process [23] (with parameters *b* = 0.0085, *δ* = 0.0028) independently in each subpopulation while allowing for dispersal events between them (with rate *γ* = 0.001). Then we varied both the cost of cooperation *c ∈* [0, 0.1*b*] and the metapopulation size (number of subpopulations) *M ∈* [1, 25] and simulated *L* = 100 independent realization for each parameter setting. We also investigated whether an explicit genotype-phenotype map and mutation between the genotypes would affect the results.

This produces a data tensor *X* with three dimensions, which can be averaged in two different ways depending on which dimension the average is taken first (see Fig. 5A). Non-ergodicity of the model manifests as the inequality of the averages (Fig. 5B). Averaging first over the subpopulations gives the ensemble average, or the metapopulation frequency

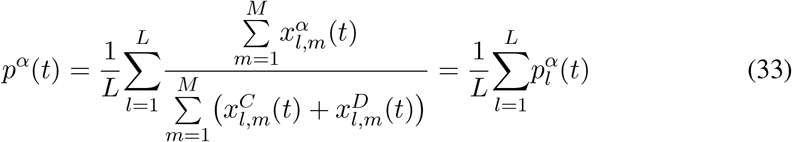

where 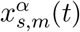 is the absolute abundance of type *α ∈ {C, D}* at time *t* in subpopulation *m* in run *s*. In contrast, averaging first over the entire length of full trajectories in time gives the typical frequency

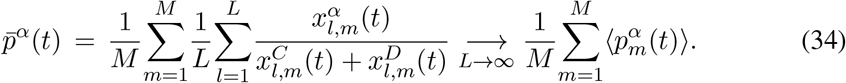

Thus, for real metapopulations with *M >* 1 and non-equal population sizes between the subpopulations (as in classical island models), the metapopulation frequency will generally differ from the average subpopulation frequency, when one is correctly computing the actual number of genotypes present in the community.

In Fig. 5C we quantified non-ergodicity using the Thirumalai-Mountain metric as

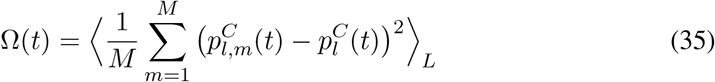

which measures how different the subpopulation frequencies are from the metapopulation frequencies averaged over all the realizations.

In Fig. 5D, we computed the total fraction of time spent in cooperator dominated 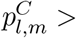 and defector dominated 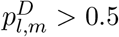 states either in absolute terms (taking into account empty states), or conditioning on occupied patches.

## Data availability

All the analysis notebooks (additional derivations) and simulation code will be made available.

## Acknowledgements

We would like to acknowledge Simo Särkkä for discussions related to the derivation of the establishment threshold.

## Author contributions

(**T.K**.) Conceptualization, Formal analysis, Investigation, Methodology, Writing – original draft, Writing – review & editing. (**A.M**) Formal analysis, Investigation, Methodology, Writing review & editing. (**V.M**.) Formal analysis, Investigation, Methodology, Supervision, Writing review & editing.

